# Translation-dependent downregulation of Cas12a mRNA by an anti-CRISPR protein

**DOI:** 10.1101/2022.11.29.518452

**Authors:** Nicole D. Marino, Alexander Talaie, Héloïse Carion, Matthew C. Johnson, Yang Zhang, Sukrit Silas, Yuping Li, Joseph Bondy-Denomy

**Affiliations:** Department of Microbiology and Immunology, University of California, San Francisco, San Francisco, CA 94158, USA; Quantitative Biosciences Institute, University of California, San Francisco, San Francisco, CA 94158, USA; Innovative Genomics Institute, Berkeley, CA 94720, USA

## Abstract

Bacteria have evolved multiple defense systems, including CRISPR-Cas, to cleave the DNA of phage and mobile genetic elements (MGE). In turn, phage have evolved anti-CRISPR (Acr) proteins that use novel and co-opted mechanisms to block DNA binding or cleavage. Here, we report that an anti-CRISPR (AcrVA2) unexpectedly inhibits Cas12a biogenesis by triggering translation-dependent destruction of its mRNA. AcrVA2 specifically clears the mRNA of Cas12a by recognizing and binding its N-terminal polypeptide. Mutating conserved N-terminal amino acids in Cas12a prevents binding and inhibition by AcrVA2 but also decreases Cas12a anti-phage activity. This mechanism therefore enables AcrVA2 to specifically inhibit divergent Cas12a orthologs while constraining its ability to escape inhibition. AcrVA2 homologs are found on diverse MGEs across numerous bacterial classes, typically in the absence of Cas12a, suggesting that this protein family may induce similar molecular outcomes against other targets. These findings reveal a new gene regulatory strategy in bacteria and create opportunities for polypeptide-specific gene regulation in prokaryotes and beyond.

## Main

Bacterial viruses (phages) are the most abundant biological entities on earth^1^. The intense selective pressure that phages impose on bacteria has spurred the evolution of many bacterial defense systems, including CRISPR-Cas^2^. In turn, phages have evolved anti-CRISPR (Acr) proteins to block CRISPR-Cas targeting^3^. Most known anti-CRISPRs inhibit cleavage by binding CRISPR-Cas complexes directly and preventing target binding or conformational activation, while others enzymatically modify the complex. The broad-spectrum inhibitor AcrVA1, for example, inactivates Cas12a by cleaving its CRISPR RNA (crRNA)^4,5^. *AcrVA1* was found encoded next to another Cas12a inhibitor, *acrVA2*, that is notably large (∼1 kb) and widely distributed across MGEs in diverse classes of bacteria^6^. AcrVA2 potently inhibits MbCas12a (*<u>M</u>oraxella <u>b</u>ovoculi*) in bacteria but not in human cells, but its mechanism has remained unclear. Here, we show that AcrVA2 recognizes the polypeptide sequence of diverse Cas12a orthologs to interrupt its biogenesis and trigger mRNA destruction independently of the promoter or codon sequence.

### AcrVA2 specifically downregulates mRNA and protein of divergent Cas12 orthologs

AcrVA2 binds MbCas12a but does not inhibit DNA cleavage *in vitro* (Extended Data Fig. 1), consistent with its inability to inhibit Cas12a in human cells^6^. This result suggested that AcrVA2 may inhibit Cas12a upstream of ribonucleoprotein (RNP) complex formation. To test this, we used strains of *Pseudomonas aeruginosa* in which plasmid-borne AcrVA2 robustly inhibits MbCas12a or LbCas12a expressed from the chromosome via different inducible promoters (Fig. 1a). Intriguingly, AcrVA2 reduced mRNA and protein levels greater than ten-fold for both MbCas12 and LbCas12a, which share 35% amino acid identity. The degree to which AcrVA2 and AcrVA2.1 (an ortholog with 84% identity) downregulated MbCas12a and LbCas12a correlated well with their ability to inhibit both orthologs (i.e. AcrVA2.1 was less active against LbCas12a). AcrVA2 downregulates codon-modified Cas12a equally well as native Cas12a (Fig. 1b,c and Extended Data Fig. 2), indicating that a specific Cas12a nucleotide sequence is not required for its downregulation. The codon-modified sequences of LbCas12a and MbCas12a were used for the rest of this study.

**Figure 1.**
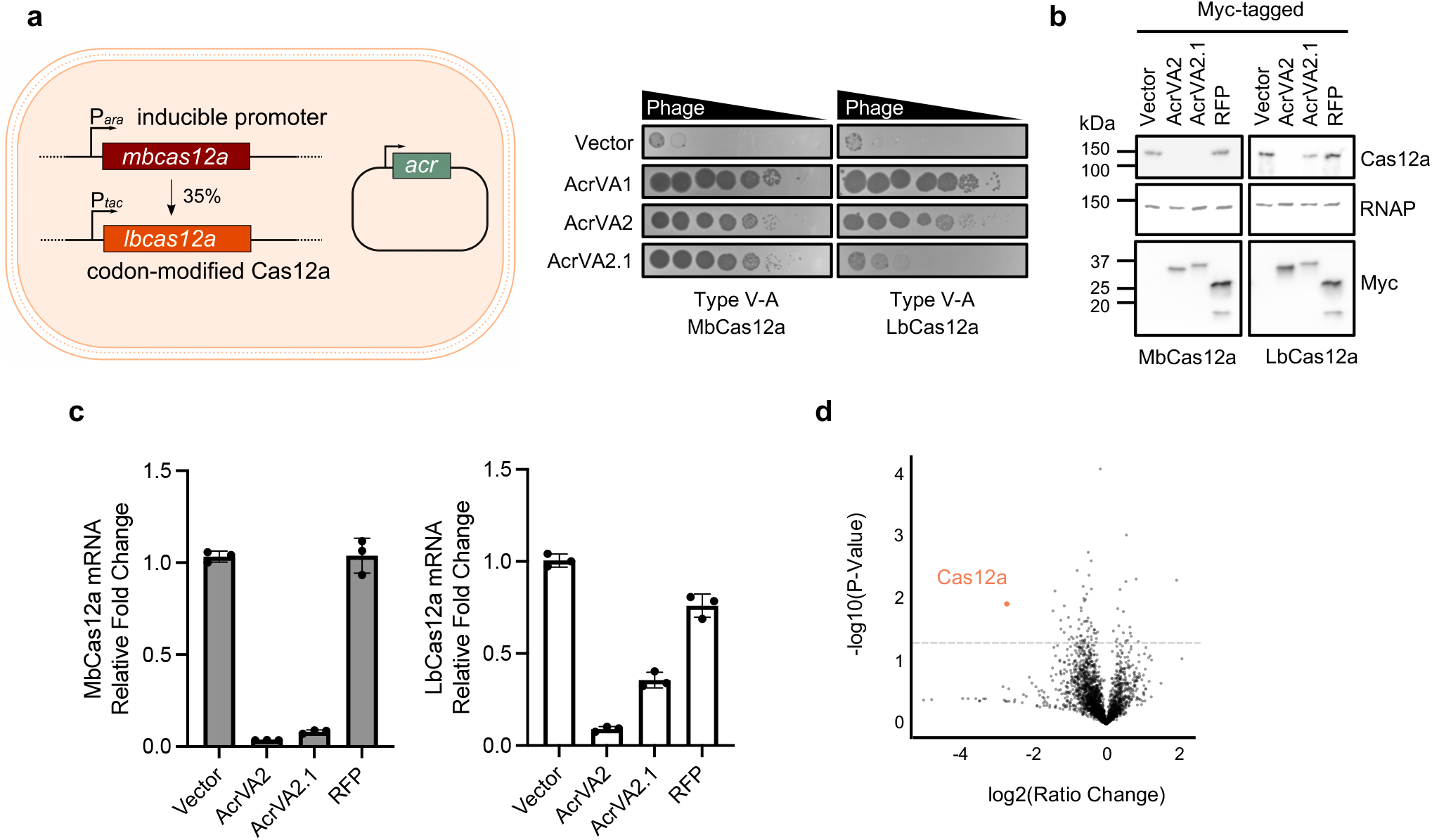
AcrVA2 specifically downregulates mRNA and protein of divergent Cas12a orthologs. (**a**) Left: Schematic of *Pseudomonas aeruginosa* strains engineered to express MbCas12a or codon-modified LbCas12a from inducible promoters and *acr* genes from plasmids. Right: Phage plaque assay with ten-fold serial dilutions of phage to assess CRISPR-Cas12a inhibition. (**b**) Western blot on bacterial lysates to assess the effect of myc-tagged AcrVA2 or control proteins on Cas12a expression. RNAP, RNA polymerase (loading control). (**c**) qRT-PCR on mRNA from bacteria expressing Cas12a and AcrVA2 or controls. Error bars indicate standard deviation. (**d**) Volcano plot for transcriptomic analysis. mRNA was extracted from bacteria expressing MbCas12a and AcrVA2 or controls. log2(Ratio Change) is the mean expression level for samples expressing AcrVA2 or AcrVA2.1 relative to controls (see methods for details). Each dot represents one gene. Dotted line indicates p-value of 0.05.

**Figure 2.**
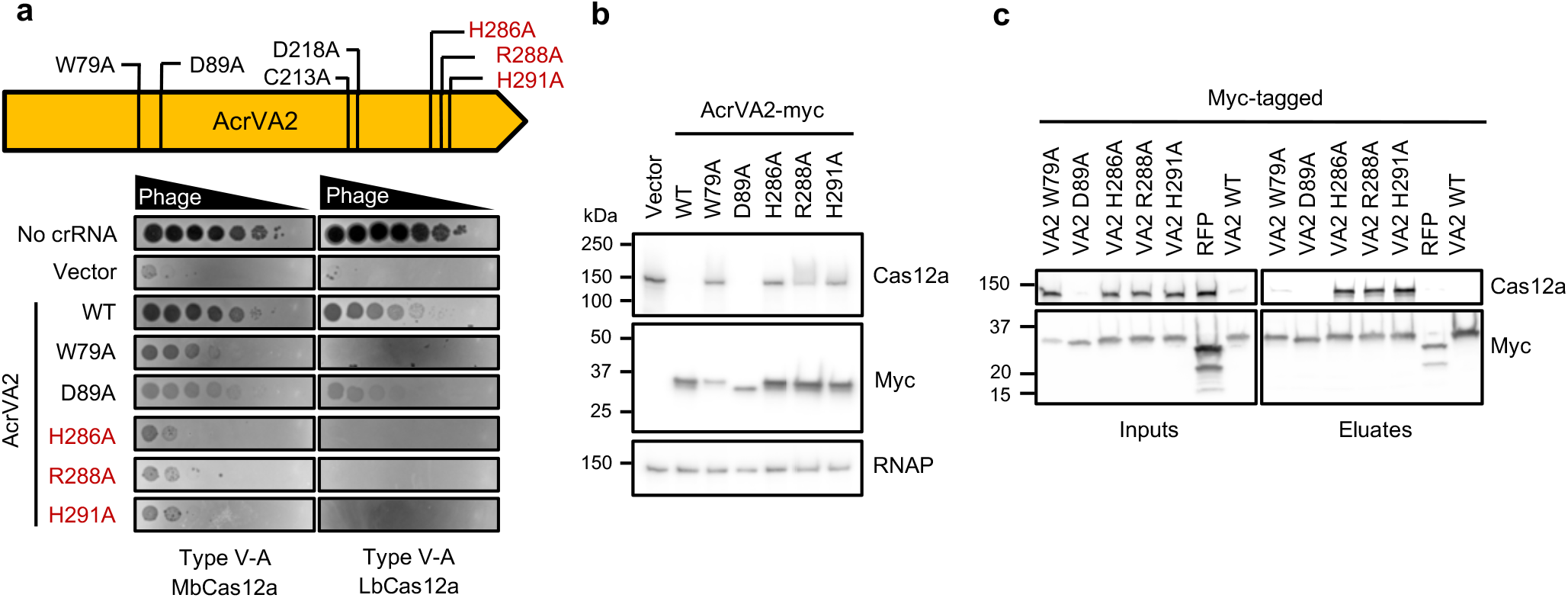
Inactive AcrVA2 mutants stably bind Cas12a protein. (**a**) Phage plaque assay using ten-fold serial dilution of phage to assess Cas12a inhibition by wildtype or mutant AcrVA2. (**b**) Western blot on bacterial lysates to assess effect of AcrVA2 point mutants on Cas12a expression. RNAP, RNA polymerase (loading control). (**c**) Immunoprecipitations on myc-tagged AcrVA2 H286A or GST control from bacterial lysates to assess interaction with Cas12a. Samples were resolved by SDS-PAGE and probed via Western blot.

The ability of AcrVA2 to downregulate divergent *cas12a* transcripts expressed from non-native promoters and featuring dramatic nucleotide changes prompted us to assess its specificity. Transcriptomic analysis revealed that *cas12a* was the only expressed gene that was significantly downregulated by AcrVA2 (Fig. 1d), while a hypothetical protein (PA3431) and a PpiC-type peptidyl-prolyl cis-trans isomerase (PA3871) were upregulated for reasons that are unclear. A closer analysis of the *cas12a* open reading frame revealed that reads were reduced by AcrVA2 at the 5’ end and were barely detectable for the latter 75% of the gene (Extended Data Fig. 3). Altogether, these results demonstrate that AcrVA2 specifically downregulates divergent and codon-modified Cas12a orthologs independently of the promoter and codon sequence.

**Figure 3.**
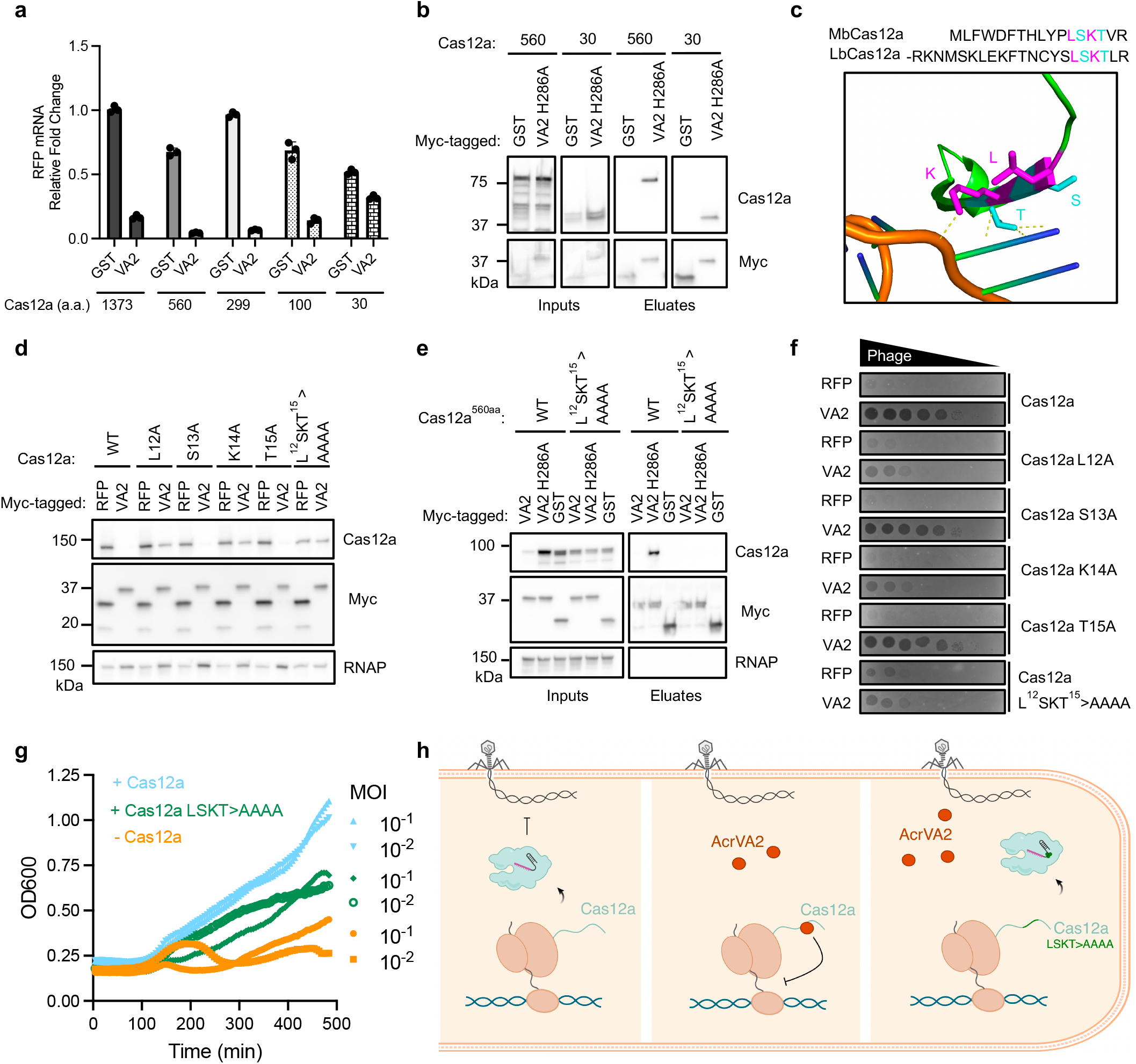
AcrVA2 binds the N-terminal polypeptide of Cas12a to trigger mRNA downregulation. (**a**) qRT-PCR on mRNA from bacteria expressing truncated Cas12a-RFP fusions to assess downregulation by AcrVA2 (VA2) relative to GST control. Numbers indicate amino acid (a.a.) length of Cas12a from N-terminus. Error bars indicate standard deviation. (**b**) Immunoprecipitations on myc-tagged AcrVA2 H286A or GST control from bacterial lysates to assess interaction with Cas12a. (**c**) Residues L^31^SKT^34^ from a crystal structure of LbCas12a with crRNA (orange sugar-phosphate helix and blue-green nucleobases)^10^. LK and ST indicated in magenta and teal, respectively. (**d**) Western blot on bacterial lysates to assess downregulation of wildtype or mutant Cas12a protein by AcrVA2 relative to RFP control. RNAP, RNA polymerase (loading control). (**e**) Immunoprecipitations on myc-tagged AcrVA2, AcrVA2 H286A or GST control from bacterial lysates to assess interaction with truncated Cas12a^560aa^. (**f**) Phage plaque assay on strains expressing wildtype or mutant Cas12a to assess inhibition by AcrVA2 relative to RFP. (**g**) Growth curves of bacteria infected with phage at different multiplicities of infection (MOI). Bacteria expressed wildtype MbCas12a (blue), MbCas12a L^12^SKT^15^>AAAA (green), or no Cas12a (orange) along with a phage-specific crRNA. (**h**) Model of AcrVA2 mechanism. Cas12a cleaves phage at specific sites to halt infection (left). AcrVA2 inhibits Cas12a biogenesis by recognizing conserved residues in its nascent polypeptide and triggering destruction of its mRNA, allowing phage infection to proceed (middle). Mutating conserved Cas12a residues prevents binding and downregulation by AcrVA2 but impairs Cas12a anti-phage function, constraining its ability to escape (right).

### Inactive AcrVA2 mutants stably bind Cas12a protein

AcrVA2 does not appear to have any conserved domains or catalytic residues that suggest a molecular mechanism for this downregulation. The crystal structure of AcrVA2 revealed three distinct domains but likewise did not resemble any known enzymes^7^. To identify amino acids in AcrVA2 that are important for its function, we mutated residues that are highly conserved across diverse AcrVA2 orthologs, including FinQ from *E. coli* ^6,8^. Interestingly, W79A and D89A mutations caused a moderate loss of function, while H286A, R288A, and H291A substitutions in the C-terminal region of the protein diminished AcrVA2 activity and restored Cas12a protein (Fig. 2a,b and Extended Data Fig. 4) and mRNA (Extended Data Fig. 3).

**Figure 4.**
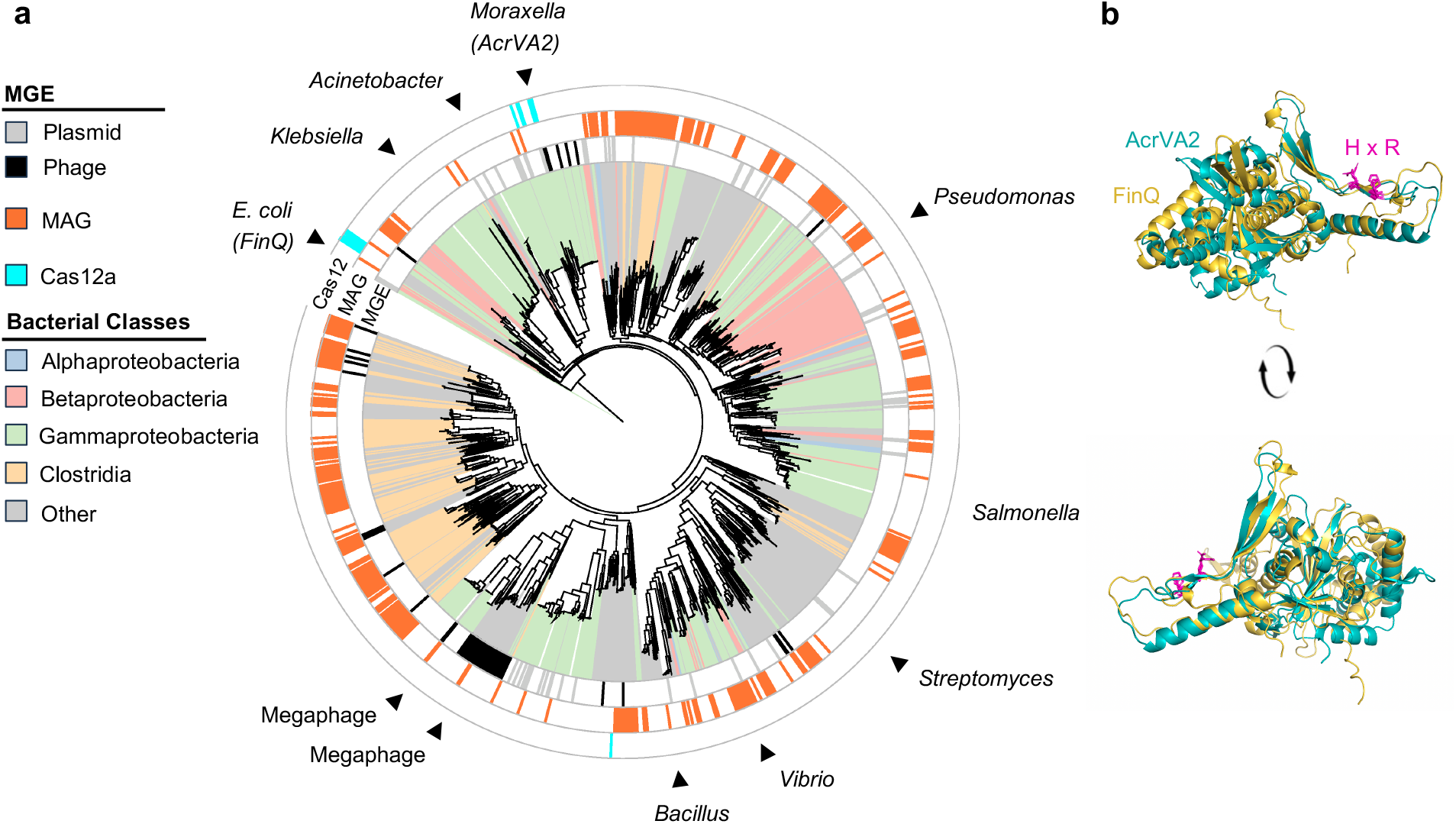
AcrVA2 orthologs are encoded on mobile genetic elements across diverse bacteria. (**a**) Approximate maximum likelihood phylogenetic tree of ∼1,800 AcrVA2 orthologs. The innermost ring indicates bacterial class by color, while the adjacent ring indicates the mobile genetic element (MGE) the ortholog is present on (where annotated). The middle ring indicates if the ortholog is found on a metagenomic assembled genome (MAG), and the outermost ring indicates if Cas12a was found in the host genome. The relative branch lengths reflect evolutionary distances between taxa without modification (i.e. tree scale is set to 1) (**b**) Alignment of predicted structures of AcrVA2 (teal) and FinQ (yellow) from Alpha Fold using Pymol. Conserved HxR motif is indicated in magenta.

The inability of AcrVA2 mutants to fully downregulate Cas12a in bacteria enabled us to ask whether these two proteins interact *in vivo*. Immunoprecipitation of myc-tagged AcrVA2 mutants (H286A, R288A, H291A) showed stable co-precipitation with Cas12a (Fig. 2c). AcrVA2^W79A^, on the other hand, abrogated downregulation of Cas12a but did not yield as robust an interaction as the C-terminal mutants, suggesting that this residue is important for the interaction with Cas12a or for AcrVA2 stability.

AcrVA2 binds apoCas12a *in vitro* (Extended Data Fig. 1a) and co-purifies with a fragment (residues 620-636) from Cas12a^7^. However, a triple mutation in AcrVA2 (E98A/D129A/D195A) that was previously shown to break this specific interaction did not significantly affect downregulation or inhibition in our *in vivo* assays (Extended Data Fig. 5), indicating that interaction at this site in Cas12a is not essential for inhibition. Overall, these results reveal that conserved C-terminal residues in AcrVA2 (H286/R288/H291) are important for mRNA downregulation and confirm that protein-protein interactions between Cas12a and AcrVA2 are not directly inhibitory.

### AcrVA2 recognizes conserved residues in the Cas12a N-terminal polypeptide

To find the region of Cas12a that is sufficient for AcrVA2 binding and downregulation, we truncated MbCas12a from the C-terminus and fused the remaining fragments to RFP. Probing RFP mRNA revealed that AcrVA2 requires only the first 100 amino acids (∼1/14th) of Cas12a to trigger mRNA downregulation (Fig. 3a). Although the first 30 amino acids of Cas12a were insufficient for downregulation, this region stably co-precipitated with AcrVA2^H286A^, suggesting that AcrVA2 recognizes and binds a sequence within this region (Fig. 3a,b). Conversely, a Cas12a mutant lacking the N-terminal 30 amino acids was well expressed but was no longer downregulated by AcrVA2 (Extended Data Fig. 6). Altogether, these data indicate that the N-terminal region of Cas12a is necessary and sufficient for AcrVA2-induced downregulation.

Comparing the N-terminal polypeptides of LbCas12a and MbCas12a revealed a conserved LSKT sequence that interacts directly with crRNA^9^ (Fig. 3c). Mutating either L12 or K14 in MbCas12a to alanine diminished downregulation, while mutating all four of these residues abolished it (Fig. 3d and Extended Data Fig. 7). As seen with codon-modified versions of Cas12a (Fig. 1b,c and Extended Data Fig. 2), synonymous mutations at this site had no effect on downregulation, showing that the amino acid sequence—rather than nucleic acid sequence—of Cas12a is the recognized substrate (Extended Data Fig. 7). Consistent with a role for the translated polypeptide, omitting the start codon from Cas12a prevented its translation and dramatically decreased mRNA downregulation by AcrVA2 (Extended Data Fig. 8).

We next assessed whether MbCas12a^LSKT>AAAA^ interacts with AcrVA2^H286A^ or wildtype AcrVA2. Because AcrVA2 binds multiple regions in MbCas12a (amino acids 1-30 and 620-636), we used truncated MbCas12a^560aa^ that lacks the PID binding site. While MbCas12a^560aa^ is downregulated by wildtype AcrVA2, co-expression with AcrVA2^H286A^ surprisingly increased MbCas12a^560aa^ protein levels, presumably due to the stabilizing effect from their interaction (Fig. 3b,e). Notably, the LSKT>AAAA mutations in the MbCas12a polypeptide sequence abolished this interaction with AcrVA2 (Fig. 3e and Extended Data Fig. 9).

Given that LSKT is recognized by AcrVA2, we next tested if these Cas12a residues are important for anti-phage function. While phage targeting by MbCas12a^LSKT>AAAA^ was only mildly defective at high expression levels (Fig. 3f), this defect became more pronounced at lower levels of induction (Fig. 3f,g and Extended Data Fig. 10). Across these conditions, MbCas12a^LSKT>AAAA^ was no longer susceptible to inhibition by AcrVA2 (Fig. 3f and Extended Data Fig. 10). Mutations in L12 and K14 also diminished targeting and susceptibility to AcrVA2, and the degree of inhibition against the different mutants correlated closely with the levels of downregulation (Fig. 3d,f). The interaction between K14/T15 and the crRNA may explain the importance of these residues for CRISPR-Cas12a activity (Fig. 3c).

Altogether, these data demonstrate that AcrVA2 recognizes and binds the Cas12a N-terminal polypeptide to drive its mRNA downregulation and inhibition (Fig. 3h). The ability to recognize conserved and functionally important residues in the target enables AcrVA2 to specifically downregulate diverse Cas12 orthologs while limiting their ability to escape inhibition.

### AcrVA2 orthologs are broadly distributed on diverse mobile genetic elements

Phylogenetic analysis revealed that AcrVA2 is unusually widespread across diverse bacterial classes and is typically found in bacteria where Cas12a is not present (Fig. 4a). We found orthologs of AcrVA2 on different types of mobile genetic elements, including plasmids and megaphages. Notably, the protein FinQ (an AcrVA2 homolog) was previously found on F-like and I-like plasmids in *E. coli*, where it was shown to inhibit conjugation of F-plasmid^11–13^. Although the mechanism of inhibition was never determined, FinQ appeared to decrease mRNA levels of F-plasmid transfer genes^13^.

Sequence alignments of divergent AcrVA2 orthologs revealed that functionally important residues in the C-terminus (H286 and R288) are conserved (Extended Data Fig. 11). The predicted structures of AcrVA2 and FinQ also show notable similarities in the C-terminal domain (Fig. 4b) and diverge near the N-terminus. These findings suggest that this regulatory paradigm may be widespread in bacteria against different targets to facilitate conflict between mobile genetic elements.

## Discussion

In this study, we have shown that AcrVA2 interrupts Cas12a biogenesis by recognizing conserved residues in its N-terminal polypeptide and triggering degradation of its mRNA. Multiple lines of evidence support this: first, AcrVA2 downregulates mRNA of divergent Cas12a orthologs independently of the promoter and codon sequence. Second, the N-terminal region of Cas12a is necessary and sufficient for this downregulation and stably binds AcrVA2. Finally, amino acid mutations (but not synonymous mutations) near the N-terminus of Cas12a abolish binding, downregulation, and inhibition by AcrVA2. The most straightforward model is that AcrVA2 recognizes the nascent polypeptide of Cas12a and triggers destruction of its mRNA before translation is complete. Although surprising, this strategy enables AcrVA2 to recognize a conserved and functionally important element in Cas12a and destroy it before it is fully expressed.

Inhibiting biogenesis is presumably ineffective against pre-existing Cas12a present in the cell. For the experiments shown here, Cas12a and AcrVA2 were induced simultaneously. However, the prophage encoding *acrVA2* in *Moraxella bovoculi* also encodes *acrVA1*, which inactivates crRNA-loaded Cas12a complexes. The dual strategies employed by these co-encoded anti-CRISPRs to inactivate crRNA-loaded Cas12a complexes (i.e. AcrVA1) and suppress Cas12a expression (i.e. AcrVA2) likely enable initial infection and stable lysogeny more effectively than either strategy alone (Extended Data Fig. 12). Dual mechanisms that inactivate complexes and reduce expression by a different mechanism have also been observed previously for phage-encoded Cas9 anti-CRISPR proteins^14^.

Some ribonucleases have previously been reported to bind the ribosomal aminoacylation (A)-site and cleave mRNAs in response to stress^15^. The nascent chain of DnaA was also shown to modulate translation elongation in response to nutrient availability^16^. To our knowledge, AcrVA2 is the first example in prokaryotes of a protein triggering mRNA degradation of a specific substrate upon recognizing its translated polypeptide sequence.

It remains unclear how AcrVA2 triggers Cas12a mRNA destruction. A similar mechanism has been demonstrated through multiple studies for tubulin autoregulation in mammalian cells^17–21^, but the mechanism for mRNA degradation has also not yet been reported. The factors and pathways involved in this fascinating mechanism will need to be elucidated in future studies.

The arms race between bacteria and phage has yielded many exciting tools and key biological discoveries for gene editing and gene regulation. Here, we show a novel strategy for CRISPR-Cas regulation that may be pervasive in microbial antagonism. The insights from this work can be applied to achieve constitutive long-term inactivation of nucleases during gene editing and enable protein-specific gene regulation in bacteria and beyond. As we explore the amazing microbial diversity in nature, many more discoveries doubtless await.

## Methods

### Bacterial strains and growth conditions

*Pseudomonas aeruginosa* strain PAO1 was used in this study. Human codon-modified MbCas12a and LbCas12a strains (Fig. 1 and Extended Data Fig. 2) were published previously^1,2^, while other strains were generated in this work (Supplementary Table 1). Strains were grown at 37 °C in lysogeny broth (LB) agar or liquid medium, which was supplemented with 50 μg ml^−1^ gentamicin, 30 μg ml^−1^ tetracycline, or 250 μg ml^−1^ carbenicillin as needed for plasmid selection or with 30 μg ml^−1^ gentamicin or 100 μg ml^−1^ carbenicillin for plasmid maintenance. MbCas12a and LbCas12a are expressed from the araBAD and tac promoters, respectively, while the crRNA is expressed from the araBAD promoter. MbCas12a-expressing strains were therefore induced with 0.3% arabinose, while LbCas12a-expressing strains were induced with 1mM isopropyl β-D-1-thiogalactopyranoside (IPTG) and 0.3% arabinose.

### Phage isolation

Phage lysates were generated by mixing 10 μl phage lysate with 150 μl overnight culture of *P. aeruginosa* and pre-adsorbing for 15 min at 37 °C. The resulting mixture was then added to molten 0.7% top agar and plated on 1% LB agar overnight at 30 °C. The phage plaques were harvested in SM buffer (100 mM NaCl, 8 mM MgSO4, 50 mM Tris-HCl, pH 7.5, 0.01% gelatin), centrifuged to pellet bacteria, treated with chloroform, and stored at 4 °C.

### Strain Engineering

Transformations of *P. aeruginosa* PAO1 strain were performed using standard electroporation protocols. Briefly, 1 ml of overnight culture was washed twice in 300 mM sucrose or 10% glycerol and concentrated tenfold. The resulting competent cells were transformed with 30 or 300 ng plasmid (for extrachromosomal uptake or chromosomal integration, respectively), incubated in antibiotic-free LB for 1 hr at 37 °C, plated on LB agar with selective media, and grown overnight at 37 °C. Chromosomal integration of pTN7C130 derivatives was achieved by co-electroporation with pTNS3, as described previously ^3,4^. For selection of pTN7C130 chromosomal integration, LB was supplemented with 30 μg ml^−1^ gentamicin. For extrachromosomal selection of pHERD30T or pHERD20T, LB was supplemented with 50 μg ml^−1^ gentamicin or 250 μg ml^−1^ carbenicillin, respectively.

### Bacteriophage plaque assays

Plaque assays were performed using 1.5% LB agar plates and 0.7% LB top agar, both of which were supplemented with 10 mM MgSO4. 150 μl overnight culture were resuspended in 3-4 ml molten top agar and plated on LB agar to create a bacterial lawn. Ten-fold serial dilutions of phage were prepared in SM phage buffer (100 mM NaCl, 8 mM MgSO4, 50 mM Tris-HCl, pH 7.5, 0.01% gelatin), spotted onto the plate, and incubated overnight at 30 °C. Agar plates were supplemented with 1mM isopropyl β-D-1-thiogalactopyranoside (IPTG) and 0.3% arabinose for assays performed with LbCas12a-expressing strains and 0.3% arabinose for assays performed with MbCas12a-expressing strains. Agar plates were supplemented with 50 μg ml−1 gentamicin or 100 μg ml−1 carbenicillin for pHERD30T and pHERD20T retention, respectively. Anti-CRISPR activity was assessed by measuring replication of the CRISPR-sensitive phage JBD30 on bacterial lawns relative to the vector control. Plate images were obtained using Gel Doc EZ Gel Documentation System (BioRad) and Image Lab (BioRad) software.

### Cloning

The native MbCas12a(237) open reading frame was amplified from genomic DNA of the 237 *Moraxella bovoculi* strain by PCR and cloned into the pTN7C130 vector using HiFi assembly (NEB). The pTN7C130 vector is a mini-Tn7 vector that integrates into the attTn7 site of *P. aeruginosa* and expresses cargo genes from the araBAD promoter.

Cas12a mutants were generated from the original vector (pTN7C130-MbCas12a, which expresses human-codon optimized MbCas12a and a C-terminal 3xHA tag)^1^ using site-directed mutagenesis. Specifically, HA-tagged constructs were generated using round-the-world PCR with non-overlapping primers encoding the desired mutations. These primers were phosphorylated using T4 polynucleotide kinase (NEB) prior to PCR, and the resulting amplicons were digested with DpnI for 1-2 h at 37 ºC to destroy the template. The products were ligated using T4 ligase (NEB) at room temperature for 1 hour or overnight at 16 ºC.

pTN7C130-Cas12a^ΔAUG^ and pTN7C130-Cas12a^Δ1-30aa^ were generated from the pTN7C130-MbCas12a vector using round-the-world PCR, as described above. Primers were designed to omit the start codon (pTN7C130-Cas12a^ΔAUG^) or the first 30 amino acids (pTN7C130-Cas12a^Δ1-30aa^) with an added start codon. Cas12a C-terminal truncations were generated by amplifying the desired sequence from the pTN7C130-MbCas12a vector and fusing to RFP with Hifi Assembly (NEB).

Anti-CRISPR (AcrIIA4, AcrVA1, AcrVA2, and AcrVA2.1) and control genes (RFP and GST) encoding a C-terminal myc tag were cloned into NcoI and HindIII sites of pHERD20T-myc and pHERD30T-myc using Gibson Assembly (NEB) or Hifi Assembly (NEB). Backbone vectors were generated by digestion at the NcoI and HindIII sites or by round-the-world PCR.

Bacterial transformations for cloning were performed using E. coli DH5α (NEB) or XL-1 blues (QB3 MacroLab) according to the manufacturer’s instructions. Plasmids were miniprepped from the resulting colonies (Zymo) and Sanger sequenced (Quintara Biosciences).

### Immunoprecipitations

PAO1 strains were grown overnight at 37°C in LB supplemented with appropriate antibiotics for plasmid retention. Induction media (50 ml per strain) was inoculated 1:100 with overnight culture and grown at 37 °C until OD600 0.5 - 1. Samples were normalized by optical density and harvested at 8,000 x g for 10 min. Pellets were stored at -80°C or immediately resuspended in 1 ml lysis buffer (50 mM Tris, pH 7.5, 250 mM NaCl, 20 mM MgCl_2_, 5% glycerol, 1% NP-40) supplemented with 0.25 mg/ml lysozyme and mini protease inhibitor cocktail (Roche). Samples were left on ice for 30 min, then sonicated twice for 10 sec at 30% amplitude at 4 °C (QSonica). Debris was spun down at 14,000 rpm for 10 minutes at 4 °C and supernatants were collected. One-tenth of the sample volume was retained for input analysis, and the remaining volume was rotated overnight at 4 °C with 45 μl of anti-C-myc magnetic beads from Cell Signaling Technology (Cat. 5698) or Pierce (Cat. 88843) to enrich for myc-tagged constructs. Beads were washed 4 × 5 min while rotating using 1 ml wash buffer (50 mM Tris, 250 mM NaCl, 20 mM MgCl_2_, 5% glycerol, 0.1% NP-40) per wash. Eluates were boiled off from the beads in 50μl of Laemmli buffer (Bio-Rad).

### Western blots

PAO1 strains were grown overnight at 37 °C in LB supplemented with appropriate antibiotics for plasmid retention. Induction media (8 ml per strain) was inoculated 1:100 with overnight culture and grown at 37 °C until OD600 0.5 - 1. Samples were normalized by optical density and harvested at 8,000 x g for 2 min. Pellets were stored at -80°C or immediately resuspended in 1 ml lysis buffer (50 mM Tris, pH 7.5, 250 mM NaCl, 20 mM MgCl_2_, 5% glycerol, 1% NP-40) supplemented with 0.25 mg/ml lysozyme and mini protease inhibitor cocktail (Roche). Samples were left on ice for 30 min, then sonicated twice for 10 sec at 20% amplitude at 4 °C (QSonica). Debris was spun down at 14,000 rpm for 10 minutes at 4 °C and supernatants were collected.

Samples (from immunoprecipitation or lysate preparation) were boiled for 10 min in Laemmli buffer supplemented with BME, separated by SDS-PAGE, and transferred to polyvinylidene difluoride (PVDF) membranes. Membranes were blocked in blocking buffer (5% milk in TBS supplemented with 2.5% Tween-20) for 1 h at room temperature and then incubated overnight at 4 °C with primary antibody in blocking buffer. Cas12a-HA was detected using horseradish peroxidase (HRP)-conjugated HA antibody (Roche) at 1:5000 dilution. LbCas12a was detected using LbCpf1 (strain ND2006) mouse monoclonal antibody (Cell Signaling Technology, #91982S) at 1:2000 dilution. Myc-tagged proteins were detected using myc-tag (9B11) mouse monoclonal antibody (Cell Signaling Technology #2276S) at 1:5000 dilution. RNA polymerase (RNAP) was detected using anti-E. coli RNA polymerase β (Biolegend, #663905) at 1:5000 dilution. Goat anti-mouse IgG secondary antibody conjugated to horseradish peroxidase (HRP) (Invitrogen, #62-6520) was used for anti-Myc, anti-LbCpf1, and anti-RNAP primary antibodies. Horseradish peroxidase (HRP) was detected using enhanced chemiluminescence (ECL) kit (Pierce). Membranes were stripped between blots by incubation in stripping buffer (Thermo Fisher) for 10-15 min then washed 2 × 5 min with PBS or TBS-T.

### Protein purification

The plasmids used for MbCas12a(33362) and AcrVA1 protein purification were published previously (8) and encode the following, in order from the N-terminus: a 10x His tag, maltose binding protein (MBP), TEV protease cleavage site, the Cas12a sequence, and a C-terminal NLS sequence for gene editing assays. Anti-CRISPR plasmids for AcrVA2, AcrVA2^H286A^, and AcrIIA4 protein purification were generated by amplifying the backbone from the AcrVA1 plasmid and cloning in other open reading frames using HiFi assembly (NEB). E. coli Rosetta2 cells freshly transformed with each plasmid were grown overnight in lysogeny broth (LB) and subcultured in LB until OD600 ∼0.5. Cells were induced with 0.4mM IPTG and grown overnight at 16 – 20 °C. Cells were harvested and resuspended in lysis buffer (20 mM Tris-HCl, pH 8 at 4°C, 150mM NaCl, 10mM imidazole, 0.5% Triton X-100, 10% glycerol, 1mM TCEP or DTT supplemented with mini complete EDTA-free protease inhibitor (Roche) and 1mM PMSF), lysed by sonication, and purified using Ni-NTA resin. The eluted proteins were cleaved with TEV protease overnight at 4°C (except for AcrVA2, which precipitated out of solution upon cleavage), and purified by size exclusion chromatography using the following buffer (20mM HEPES pH 7.5, 150mM KCl, 20mM MgCl2, 10% glycerol, 1mM DTT).

### Binding assays

MBP-tagged anti-CRISPR proteins (or control proteins lacking MBP) were incubated with amylose resin at room temperature for 30 min with occasional shaking. TEV-cleaved MbCas12a(33362) protein was added and incubated for another 30 min with occasional shaking. Samples were spun at 600 x g for 2 minutes to collect the flow through. Beads were washed five times with binding buffer (20 mM Tris, pH 7.5, 200 mM NaCl) and eluted with binding buffer supplemented with 40 mM maltose. Inputs and eluates were resolved by SDS-polyacrylamide gel electrophoresis (SDS-PAGE) and stained with Coomassie Blue.

### Cleavage assays *in vitro*

In vitro Cas12a cleavage assays were performed using purified, TEV-cleaved MbCas12a^33362^ protein and crRNAs that were synthesized commercially (IDT). dsDNA templates were amplified from plasmids and purified using DNA Clean and Concentrator Kit (Zymo). Ribonucleoprotein (RNP) formation: crRNA was diluted to 500 nM in 1x cleavage buffer (20 mM HEPES-HCl, pH 7.5, 150 mM KCl, 10 mM MgCl_2_, 0.5 mM TCEP), heated at 70 ºC for 5 min, then allowed to cool down to room temperature. crRNA was mixed with 500 nM Cas12a protein at 1:1 ratio (250 mM final), then incubated at 37 ºC for 10 min. Linear template and purified Acr protein were diluted separately in cleavage buffer to 5 nM and 250 nM, respectively, and heated for 10 min at 37 ºC. Pre-formed RNPs and Acr proteins (each 25 nM final concentration) were added to template DNA and incubated at 37 ºC for 30 min. Reactions were quenched with 6x Quench Buffer (30% glycerol, 1.2% SDS, 250 mM EDTA). Reactions were loaded onto 1% TAE agarose gels and resolved by electrophoresis. Gels were post-stained with SYBR Gold (Invitrogen) for 1 hr at room temperature and imaged using BioRad Gel Doc EZ Imager.

### RNA extraction and qRT-PCR

Induction media (8 ml per strain) was inoculated 1:100 with overnight cultures and allowed to grow to OD600 of 0.5 - 1. Bacteria were harvested and resuspended in 800 μl water and mixed 1:1 with pre-heated lysis buffer (100 μl of 8x lysis solution [0.3 M sodium acetate, 8% sodium dodecyl sulfate, 16 mM EDTA] mixed with 700 μl acid phenol:chloroform per sample and heated at 65 ºC for 15 min). The lysate was then incubated at 65 ºC for 5 – 10 minutes with frequent vortexing. Samples were spun at 12,000 x g for 15 minutes at 4 ºC and the upper aqueous layer was collected. Samples were extracted two times with equivalent volumes of chloroform and incubated overnight at -20 ºC with 3 volumes of 100% ethanol. Samples were harvested at 14,000 rpm for 15 min at 4 ºC and washed with 75% ethanol. Pellets were resuspended in Milli-Q water and assessed for yield and purity using a spectrophotometer.

Purified RNA was treated for contaminating DNA using the Turbo DNase kit (Invitrogen). Relative mRNA levels of Cas12a, RFP, and RpoD control were assessed using the Luna Universal One-Step qRT-PCR kit (New England Biolabs) according to the manufacturer’s instructions. Primers were assessed for efficiency of amplification and controls lacking reverse transcriptase were used in all experiments. RpoD was used as a loading control to normalize expression. Experiments were repeated at least three times (from RNA extraction to qRT-PCR analysis) on randomly chosen colonies (typically from different transformations).

### RNAseq and analysis

RNA was extracted from bacterial strains expressing MbCas12a and either AcrVA1, AcrVA2, AcrVA2.1, AcrVA2^H286A^, or AcrIIA4 according to the protocol described above. Purified RNA was treated for contaminating DNA using Turbo DNase (Invitrogen) for 30 min at 37 ºC and then purified by phenol-chloroform extraction. RNA was prepared for sequencing using the SMARTER Stranded RNA sequencing kit (Takara Bio) according to the manufacturer’s instructions with the following parameters. Fragmentation was performed for 4 min, and PCR amplification of the final library was carried out for 10 cycles. Amplified libraries were quantified using a Qubit 4.0 fluorometer (Life Technologies) and sequenced on an Illumina HiSeq in single end format (50 bp) with one 6 bp index read.

Sequencing adapters were trimmed from the reads at the 3’ end, and 3 bp were additionally trimmed from the 5’ end of every read to account for the template switching activity of the RT using cutadapt. Trimmed reads were mapped to the PAO1 reference genome and to the Cas12a CDS using bowtie2. RPKM values were then calculated per gene and mappings were visualized using IGV.

The volcano plot was generated by first normalizing RPKM counts against the housekeeping gene rpoD. Genes for which the sum of the normalized counts were less than 10% of *rpoD* were considered low expression and were removed from further analysis. The ratio change between the experimental group (AcrVA2 and AcrVA2.1) and control group (AcrIIA4, AcrVA1, and AcrVA2^H286A^) were calculated for each gene from the normalized counts, and the p-value was determined by t-Test.

### Phylogenetic analysis

The sequence of AcrVA2 from *Moraxella bovoculi* 58069 was used to query the NCBI non-redundant protein database with PSI-BLAST (3 rounds, default parameters)^5^. Hits were filtered to remove any with a sequence length above 2 standard deviations from the mean. MAFFT alignment was used to align the roughly 1.8k homologs that remained. Because conservation is a limited domain, L-INS-i iterative refinement method was used with a maxiterate 1000 set^6^.

FastTree2 was used to compute an approximate maximum-likelihood phylogenetic tree. Available metadata associated with the proteins were used to classify hits as plasmid, phage or MAG. Proteins without associated metadata were included but not classified. Visualization was done with iTol and custom scripts were used to generate additional display features^7^. Sequences used to make the tree can be found in Supplementary Table 2.

### Protein Alignment

The protein sequence of different orthologues of AcrVA2 were aligned with Clustal Omega and colored in Jalview using BLOSUM62 scheme. Protein sequences can be found in Supplementary Table 3.

## Data availability

RNAseq data will be deposited to the Sequence Read Archive prior to publication. All other data supporting the findings in the Article and the Supplementary Information are available from the corresponding authors on request.

## Acknowledgements

We thank the Bondy-Denomy lab as well as Carol Gross (UCSF) and her lab for thoughtful discussions. We thank Kyle Watters for the pKEW-AcrVA2 plasmid. We also thank Magda Paredes and Danh Le for preparing the media used in this work.

## Author contributions

Conceptualization: NDM and JBD. Data curation: NDM, AT, HC, YZ, SS, MCJ. Formal analysis: NDM, SS, YL, MCJ. Funding acquisition: NDM and JBD. Investigation: NDM, AT, HC, YZ. Methodology: NM, SS, MCJ, JBD. Project administration: NDM, JBD. Resources: NM, AT, YZ. Supervision: NM, JBD. Validation: NDM, AT, HC, YZ. Visualization: NDM, MCJ, JBD. Writing - original draft: NDM, JBD. Writing - review & editing: NDM, HC, AT, SS, YL, and JBD.

## Competing interests

UCSF has filed a patent on the use of inhibitors of CRISPR-Cas12a, on which NDM and JBD are listed as inventors. J.B.-D. is a scientific advisory board member of SNIPR Biome and Excision Biotherapeutics, a consultant to LeapFrog Bio, and a scientific advisory board member and co-founder of Acrigen Biosciences. The Bondy-Denomy lab received research support from Felix Biotechnology.

**Correspondence and requests for materials** should be addressed to Nicole D. Marino or Joseph Bondy-Denomy.

## Additional information

### Supplementary Information

Supplementary Tables 1-3 are included with this submission.

