## Extended Data for "Translation-dependent downregulation of Cas12a mRNA by an anti-CRISPR protein"

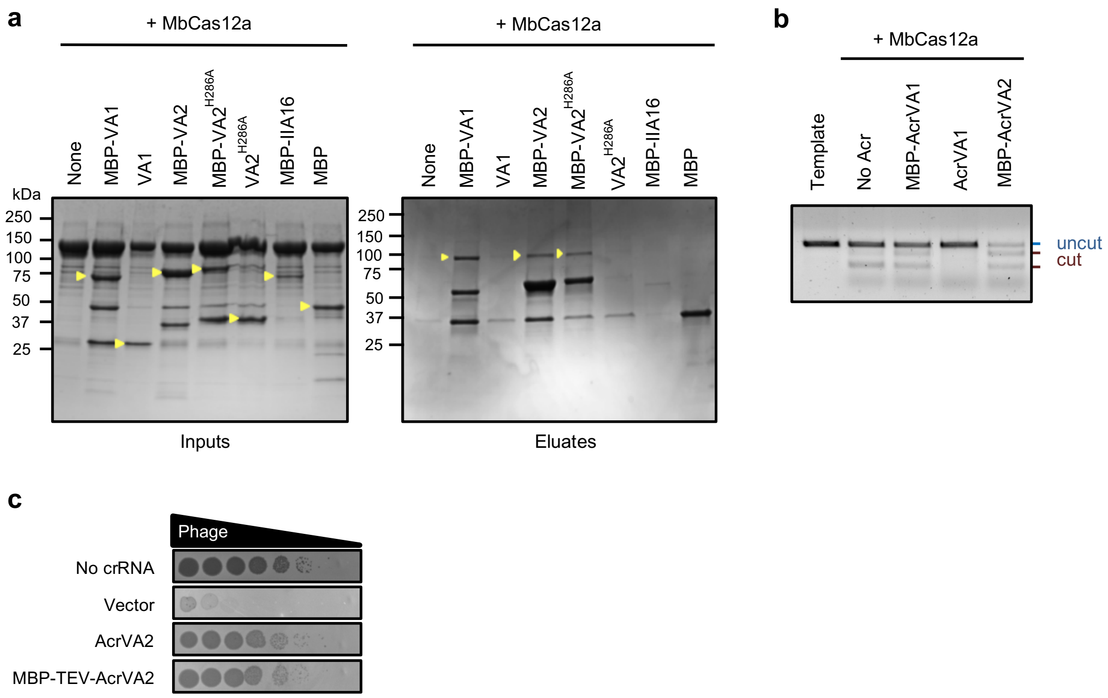


**Extended Data Fig. 1. AcrVA2 binds apoCas12a *in vitro* but does not inhibit cleavage.** (**a**) SDS-PAGE and Coomassie stain to assess anti-CRISPR binding to Cas12a. (**b**) *In vitro* cleavage assay for MbCas12a[33362] ribonucleoproteins (RNPs) to assess inhibition by anti-CRISPR proteins or control. Blue line indicates uncut template, and brown lines indicate cut template. (**c**) Phage plaque assay to test Cas12a inhibition by tagged AcrVA2 compared to untagged AcrVA2. Ten-fold serial dilutions of phage were plated on bacterial lawns to assess Cas12a inhibition.

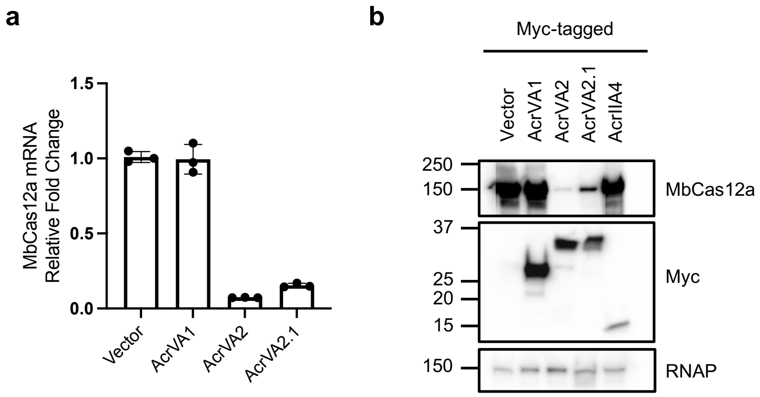


**Extended Data Fig. 2. AcrVA2 downregulates codon-modified MbCas12a.** (**a**) qRT-PCR on mRNA extracted from bacteria to assess downregulation of codon-modified Cas12a by AcrVA2 relative to controls. (**b**) Western blot on bacterial lysates to assess downregulation of codon-modified MbCas12a by AcrVA2 relative to controls. RNAP, RNA polymerase (loading control).

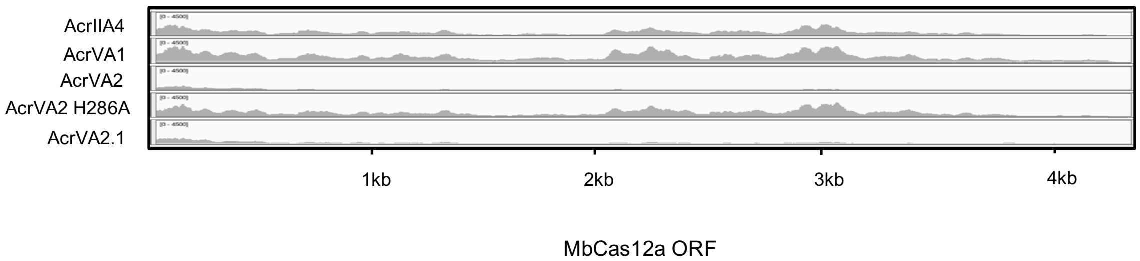


**Extended Data Fig. 3. IGV snapshot of raw RNA sequencing reads mapped to the MbCas12a open reading frame.** RPKM values for MbCas12a are as indicated for each sample: AcrIIA4, 217; AcrVA1, 316; AcrVA2, 38; AcrVA2 H286A, 225; AcrVA2.1, 71.

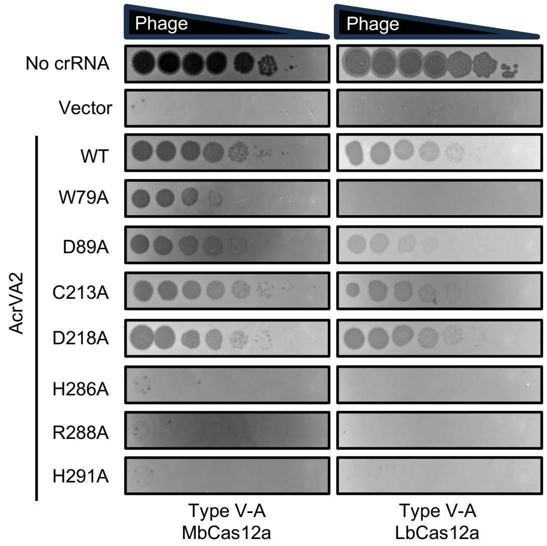


**Extended Data Fig. 4. Mutations in conserved C-terminal AcrVA2 residues abrogate Cas12a inhibition.** Phage plaque assay using ten-fold serial dilution of phage to assess Cas12a inhibition by wildtype or mutant AcrVA2.

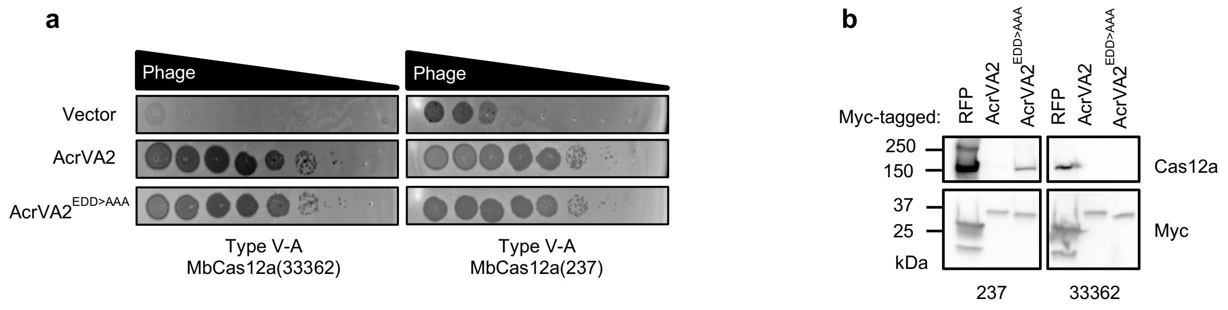


**Extended Data Fig. 5. AcrVA2 triple mutants defective in *in vitro* binding still inhibit and downregulate MbCas12a.** (**a**) Phage plaque assay on strains expressing different orthologs of MbCas12a. Ten-fold serial dilutions of phage were plated on bacterial lawns to assess Cas12a inhibition. (**b**) Western blot on bacterial lysates to assess downregulation of different MbCas12 orthologs (from strains 237 and 33362) by AcrVA2 E98A/D129A/D195A relative to controls.

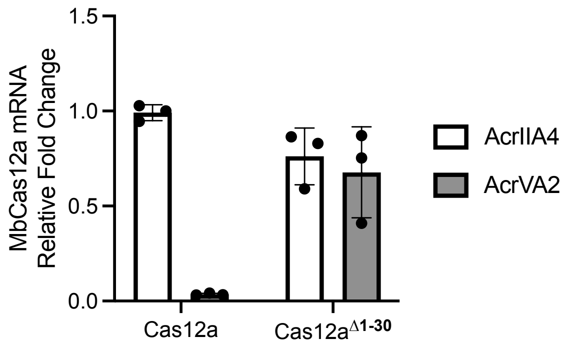


**Extended Data Fig. 6. N-terminal deletion in Cas12a abrogates downregulation by AcrVA2.** qRT-PCR on mRNA from bacteria expressing wildtype MbCas12a or MbCas12a lacking its first 30 amino acids (and provided with a start codon). Error bars indicate standard deviation.

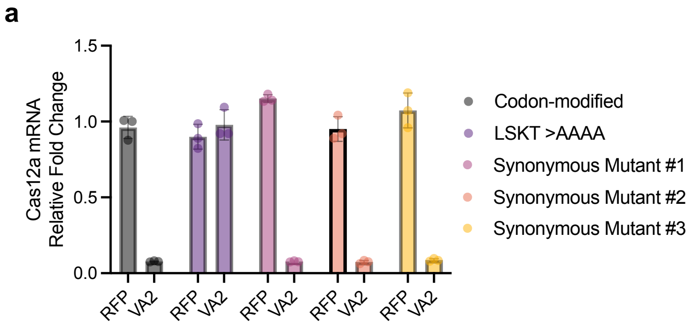


**b**

```
                                                 L12SKT15
Native             ATGTTGTTTCAAGACTTTACTCATCTATACCCCTTATCAAAAACCGTG
Codon-modified     ATGCTGTTCCAGGACTTTACCCACCTGTATCCACTGTCCAAGACAGTG
Silent mutation 1  ATGCTGTTCCAGGACTTTACCCACCTGTATCCACTCTCAAAAACGGTG
Silent mutation 2  ATGCTGTTCCAGGACTTTACCCACCTGTATCCACTCTCGAACACCGTG
Silent mutation 3  ATGCTGTTCCAGGACTTTACCCACCTGTATCCACTCTCGAACACCGTG
                   *** **** ** ******** ** ** ** **  * ** ** ** ***
```

**Extended Data Fig. 7. Non-synonymous mutations in MbCas12a L$^{12}$SKT$^{15}$ motif abolish mRNA downregulation by AcrVA2, but synonymous mutations do not.** (**a**) qRT-PCR on mRNA to assess downregulation of different Cas12a variants (with synonymous or non-synonymous codon changes) by AcrVA2 relative to RFP control. (**b**) Clustal omega alignment of nucleotide sequences for first 16 amino acids of different MbCas12a variants. Red line indicates nucleotides within L$^{12}$SKT$^{15}$ amino acid residues.

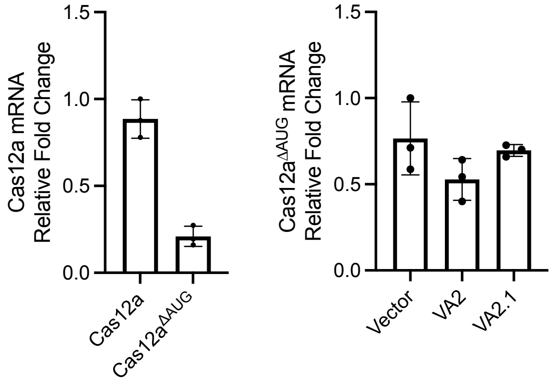


**Extended Data Fig. 8. Omitting the start codon of MbCas12a abrogates mRNA downregulation by AcrVA2 and AcrVA2.1.** (**a**) qRT-PCR on mRNA from bacteria expressing wildtype MbCas12a or MbCas12a lacking a start codon (MbCas12a$^{\Delta AUG}$). The start codon mutant of MbCas12a exhibited lower mRNA, likely due to increased exposure to ribonucleases in the cell. (**b**) qRT-PCR on mRNA from bacteria expressing MbCas12a lacking a start codon (MbCas12a$^{\Delta AUG}$) and AcrVA or control. Error bars indicate standard deviation.

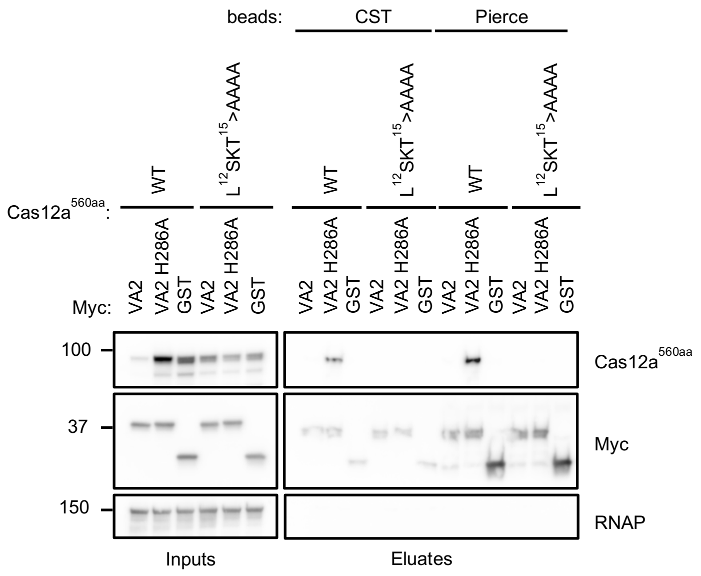


**Extended Data Fig. 9. Mutations in the first 15 amino acids of MbCas12a abrogate binding, downregulation, and inhibition by AcrVA2 (extended from Figure 4).** Immunoprecipitations on myc-tagged AcrVA2 or GST control from bacterial lysates using two different anti-myc tag magnetic beads (from CST and Pierce). Samples were resolved by SDS-PAGE and probed via Western blot. RNAP, RNA polymerase (loading control).

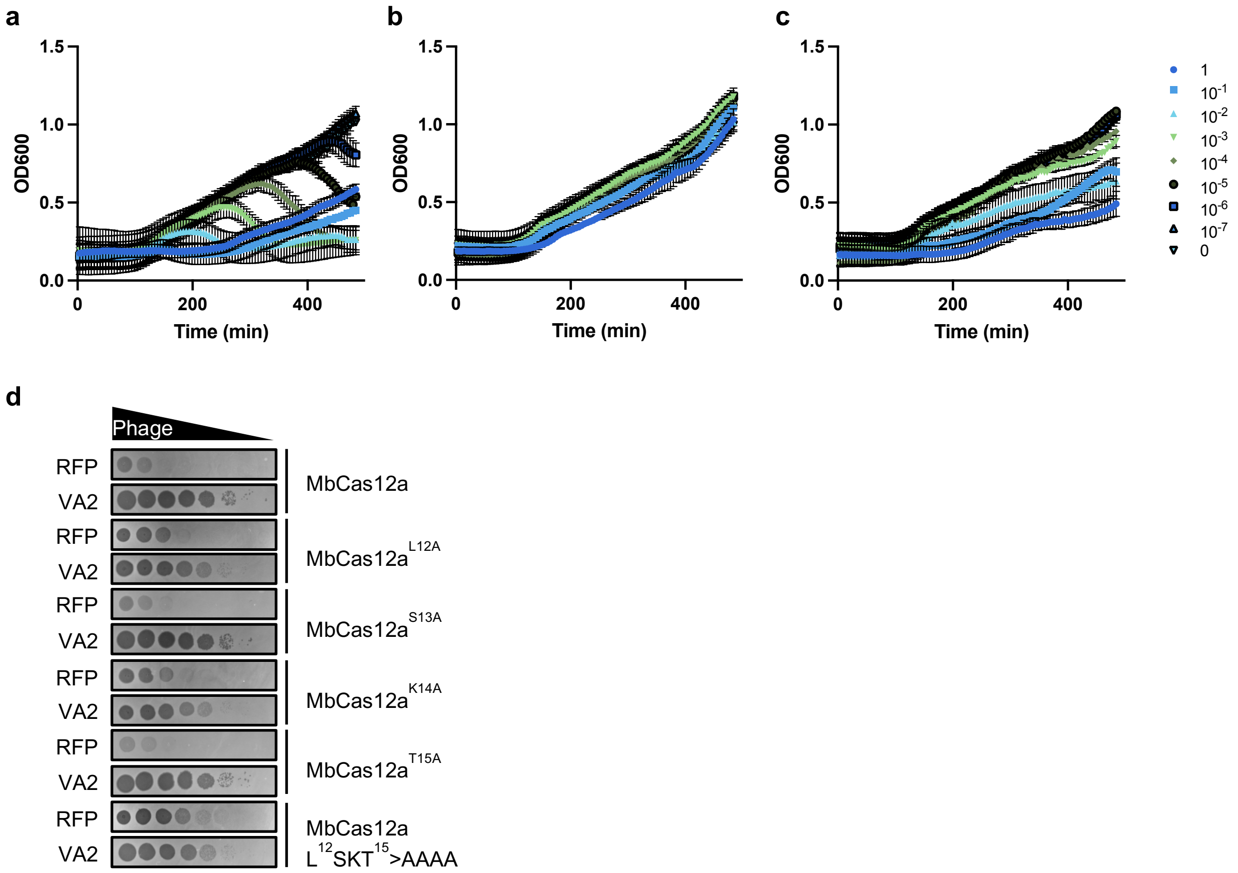


**Extended Data Fig. 10. Mutations in LSKT motif decrease Cas12a anti-phage activity.** Eight-hour growth curves of *Pseudomonas aeruginosa* PAO1 expressing (**a**) no Cas12a, (**b**) wildtype Cas12a, or (**c**) Cas12a L$^{12}$SKT$^{15}$>AAAA. Cultures were infected with phage at multiplicities of infection (MOI) indicated in the legend. Error bars indicate standard deviation. (**d**) Phage plaque assay on bacteria expressing wildtype or mutant Cas12a. Ten-fold serial dilutions of phage were plated on bacterial lawns to assess Cas12a inhibition. In this experiment, 0.1% arabinose was used to induce Cas12a, crRNA, and AcrVA2 or RFP expression.

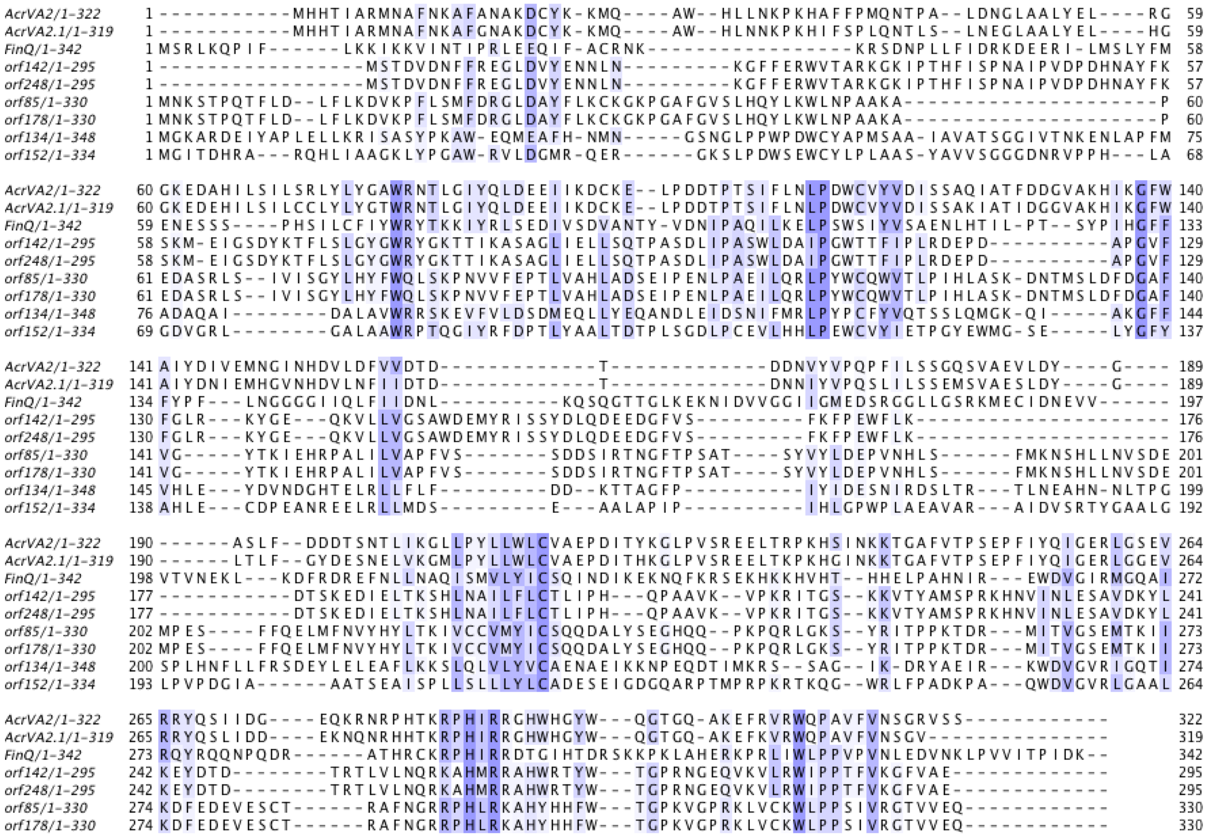


**Extended Data Fig. 11. Protein sequence alignments of AcrVA2 orthologs from *Moraxella bovoculi*, *E. coli* (FinQ), and megaphages.** Dark blue indicates that the residue matches the consensus sequence at that position, while light blue indicates that the two residues have a positive Blosum62 score. ORFs 142, 248, 85, and 178 are encoded in megaphage genomes.

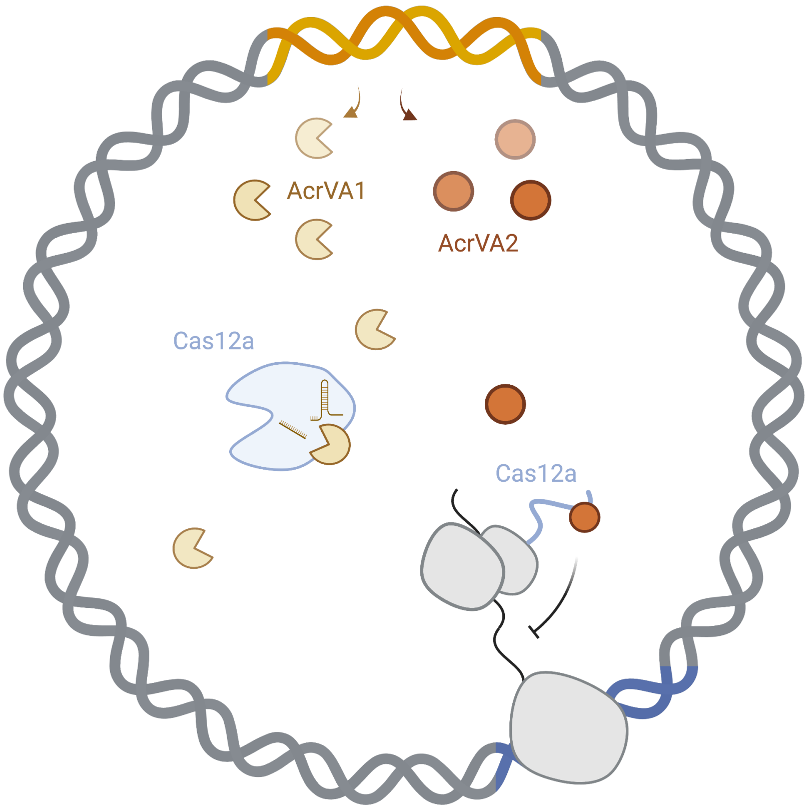


**Extended Data Fig. 12. Model of Cas12a inhibition by AcrVA1 and AcrVA2.** AcrVA1 and AcrVA2 expressed from a prophage in *Moraxella bovoculi* inhibit Cas12a from targeting the prophage genome (orange) by cleaving the crRNA and inhibiting Cas12a biogenesis, respectively.
